# Residual Partial Least Squares Learning: Brain Cortical Thickness Simultaneously Predicts Eight Non-pairwise-correlated Behavioural and Disease Outcomes in Alzheimer’s Disease

**DOI:** 10.1101/2024.03.11.584383

**Authors:** Duy-Thanh Vu, Duy-Cat Can, Christelle Schneuwly Diaz, Julien S. Bodelet, Guillaume E. Blanc, Huy Phan, Gilles Allali, Viet-Dung Nguyen, Hengyi Cao, Xingru He, Yannick Müller, Bangdong Zhi, Haochang Shou, Haoyu Zhang, Wei He, Xiaojun Wang, Marcus Munafò, Guy Nagels, Philippe Ryvlin, Nguyen Linh Trung, Giuseppe Pantaleo, Oliver Y. Chén, the Open Access Series of Imaging Studies (OASIS), the Alzheimer’s Disease Neuroimaging Initiative (ADNI)

**Affiliations:** Platform of Bioinformatics, Lausanne University Hospital, Lausanne, Switzerland; Faculty of Biology and Medicine, University of Lausanne, Lausanne, Switzerland; Department of Electrical Engineering, KU Leuven, Leuven, Belgium; Centre Leenaards de la mémoire, CHUV, Lausanne, Switzerland; École Nationale Supérieure de Techniques Avancées de Bretagne, Bretagne, France; VNU University of Engineering and Technology, Hanoi, Vietnam; Feinstein Institutes for Medical Research, Manhasset, NY, USA; Division of Psychiatry Research, Zucker Hillside Hospital, Glen Oaks, NY, USA; School of Public Health, He University, Shengyang, China; Innovation and Healthcare Group, University of Bristol, Bristol, UK; Department of Biostatistics, University of Pennsylvania, Philadelphia, PA, USA; Division of Cancer Epidemiology and Genetics, NIH, Bethesda, MD, USA; School of Psychological Science, University of Bristol, Bristol, UK; Department of Neurology, Universitair Ziekenhuis Brussel, Jette, Belgium; Institute of Biomedical Engineering, University of Oxford, Oxford, UK; Département des neurosciences cliniques, CHUV, Lausanne, Switzerland

## Abstract

Alzheimer’s Disease (AD) is the leading cause of dementia, affecting brain structure, function, cognition, and behaviour. While previous studies have linked brain regions to univariate outcomes (*e*.*g*., disease status), the relationship between brain-wide changes and multiple disease and behavioural outcomes of AD is still not well understood. Here, we propose Residual Partial Least Squares (re-PLS) Learning, an explainable and generalisable framework that models high-dimensional brain features and multivariate outcomes, accounting for confounders. Using re-PLS, we map the *many-to-many* pathways between cortical thickness and multivariate AD outcomes; identify neural biomarkers that simultaneously predict multiple outcomes; control for confounding variables; conduct longitudinal AD prediction; and perform cross-cohort AD prediction. To evaluate its efficacy, we first carry out within-cohort cross-subject validation using ADNI data, and further examine its reproducibility via between-cohort cross-validation using ADNI and OASIS data. Together, our results unveil brain regions jointly but differentially predictive of distinctive cognitive-behavioural scores in AD.

## 1 Main

Alzheimer’s disease (AD) is a neurodegenerative disorder affecting 50 million people worldwide and is projected to affect as many as 152 million by 2050 [1]. It is the most common form of dementia [2]. An early symptom of AD is difficulty remembering recent events. Gradually, a patient may exhibit language and orientation problems, mood swings, loss of motivation, self-neglect, and behavioural changes. In general, one observes progressive cognitive decline in AD, accompanied by a gradual loss of bodily functions, eventually leading to death [3]. An AD patient’s typical life expectancy following diagnosis ranges from three to nine years [4].

Discovering biomarkers associated with AD is essential in understanding the pathology of the disease, identifying patients, assessing disease progression, and enabling the timely management of the condition [5]. An important biomarker of AD is brain cortical thickness, also known as the AD cortical “signature” [6]. Changes of cortical thickness are differentially expressed across brain areas and vary between pre-clinical dementia stages (*i*.*e*., subjects with mild cognitive impairment (MCI)) and dementia [7, 8]. In general, compared to cognitively normal (CN) subjects, individuals with MCI and AD have decreased cortical thickness in the medial temporal lobe region and parts of the frontal and parietal cortices [7–9]. As the disease progresses, cortical thinning is observed across the entire cortex, especially in the lateral temporal lobe [7]. In parallel, cortical thickness of frontal, parietal, and temporal lobes in AD is correlated with cognitive impairment [8], while regional thinning predicts (even mild) AD [10].

In addition to cortical thickness changes, AD is accompanied by multiple cognitive and behavioural disruptions in memory, language, orientation, judgment, or problem-solving [11]. Yet, despite advances in single outcome assessment and prediction, our understanding of the many-to-many (*i*.*e*., many brain areas to many outcomes) relationship between the spatially varying cortical thickness changes and multiple symptoms or cognitive (dys)functions has remained limited.

To improve our knowledge about and better manage the disease, it is crucial to identify and isolate brain regions, each of whose cortical thickness may be differentially associated with a unique, or several, cognitive or behavioural outcomes, chart the pathways between each set of brain areas and their corresponding outcome, as well as quantify the pathway effect. Equally important is to leverage these pathways and parameters of the identified regions to predict multiple, likely non-pairwise-correlated, cognitive and behavioural scores. That is, one uses cortical thickness data from identified, potentially different brain regions to predict each corresponding outcome. Such quests for neurobiological insights and predictive performances require joint effort, integrating methodological innovations and biological knowledge. **First**, there is a need to search for subsets of brain areas respectively associated with different cognitive and behavioural outcomes. This is important for improving our understanding of disease pathology and aiding in pathway estimation. **Second**, there is a need to chart pathways between high-dimensional cortical thickness and multiple cognitive and behavioural outcomes. Cortical thickness changes in AD occur across several functional brain regions, each likely projecting to multiple cognitive and behavioural domains. Therefore, understanding these pathways can provide insights into how cortical thickness in different brain areas may be linked to corresponding cognitive-behavioural outcomes. **Third**, there is a need to deal with confounding variables that affect both brain data and behaviour. Indeed, cortical thickness and disease outcomes differ across age and gender groups (see Fig. 1a-d); ignoring them or only considering their association with outcomes, but not with cortical thickness features, may bias estimated pathways [12]. In disease analysis and prediction, neglecting confounding effects may yield clinical misinterpretations [13]. **Fourth**, there is a need to predict multivariate, non-pairwise-correlated outcomes. Although predictive models built for assessing single outcomes [14] have considerably advanced our understanding of general aspects (such as disease status [15]) or specific subdomains (such as cognitive decline [16]) of AD, single-outcome prediction [17] may not capture multi-dimensional and -functional cognitive and behaviour degenerative landscapes of the disease. **Fifth**, as a neurodegenerative disease that not only progresses differentially along various cognitive and behavioural domains but also develops in time, there is a need to predict AD progression longitudinally. This may help evaluate or anticipate the cognitive decline and disease conversion early and manage the disease advancement in a timely manner. **Finally**, while selected features and predictive models facilitate biological interpretation and disease assessment, to introduce them in broader practices and to endorse their scientific efficacy, there is a need to demonstrate that properties of features and predictive models can be generalised to different subjects and, particularly, reproduced in other cohorts and datasets.

**Fig. 1:**
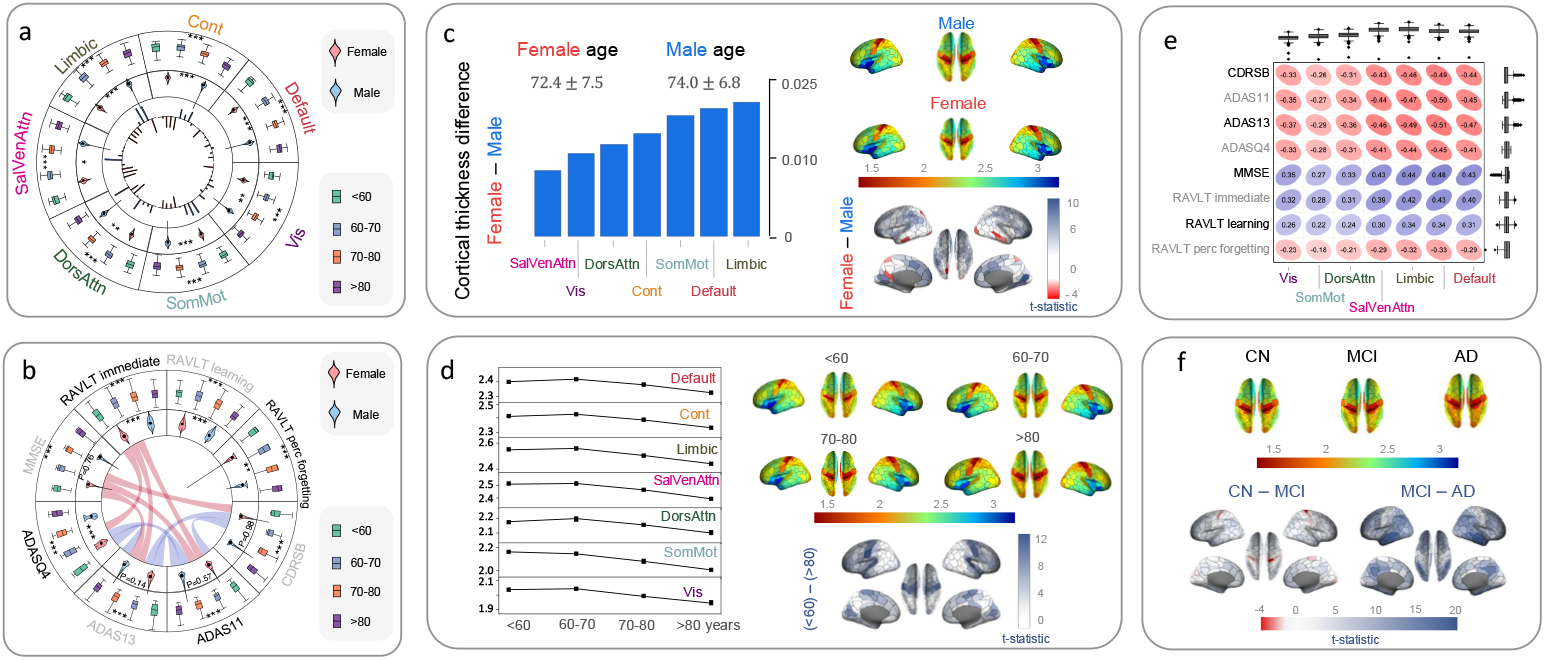
Effect of age and gender on cortical thickness as well as disease and behavioural outcomes in AD. (a) Cortical thickness from seven functional brain areas exhibits different variations across age and gender groups. Outer, middle, and inner circles show cortical thickness by age, gender, and predictive weights for eight outcomes, respectively. Outer bars represent positive weights, and inner bars represent negative weights based on averaged linear regression coefficients. Weights are normalised to [- 1, 1] within each functional network. (b) Eight AD-related outcomes vary by age and gender and are not strongly pairwise correlated. Outer, middle, and inner circles display outcomes by age, gender, and correlation structure, respectively. Connected lines indicate high correlations (|*r*| ≥ 0.7). (c) Cortical thickness exhibits gender differences across brain networks. The left plot shows mean cortical thickness differences between females and males across seven functional brain networks. The right plots show spatial distribution of gender differences across brain regions. The female cortex is generally thicker than the male cortex. (d) Cortical thickness varies by age group across brain networks. The left plot shows mean thickness decline with age across seven functional brain networks. The right plots show spatial distribution of age-related thickness changes across brain regions, with younger groups (*<* 60) generally showing thicker cortex than older groups (*>* 80). (e) Distributional and associative analysis between AD-related outcomes and regional cortical thickness. Top boxplots show cortical thickness distributions across seven functional brain areas; right boxplots show cognitive and behavioural score distributions from eight tests. The value in each ellipse represents correlations between each outcome and the corresponding brain network thickness. (f) Cortical thickness differences across diagnostic groups (CN, MCI, AD) with statistical comparisons. The top plots show mean thickness across brain regions for each diagnostic group. Bottom plots show statistical comparisons between groups (CN-MCI and MCI-AD). For panels (c), (d), and (f), rainbow colour bars indicate normalised cortical cohort-specifis; red-blue colour bars represent t-statistics for brain regions.

Here, we introduce *Residual Partial Least Squares* (re-PLS), by integrating residual learning [18, 19], partial least squares (PLS) [20–22], and predictive modelling [14, 23], to identify brain regions whose cortical thickness is associated with and predictive of multivariate, non-pairwise-correlated outcomes in AD; uncover multivariate many-to-many pathways from these regions to disease and behavioural outcomes; and predict such outcomes at both population and individual levels, across cross-sectional and longitudinal settings. Specifically, we apply re-PLS to data from the Alzheimer’s Disease Neuroimaging Initiative (ADNI) and discover potential pathways between cortical thickness data and multivariate disease and behavioural outcomes while controlling for confounding age and gender variables. Furthermore, we use re-PLS to perform longitudinal AD prediction. Finally, we test the features selected from and the model trained using ADNI data, without further modelling, on data from the Open Access Series of Imaging Studies (OASIS), and *vice versa*.

## 2 Results

We begin by summarising five key points regarding our findings. **(1)** Both re-PLS and other baseline models suggest that brain cortical thickness predicts multiple, non-pairwise-correlated behavioural and disease outcomes in AD (see Fig. 3). **(2)** The re-PLS and PLS yield higher predictive accuracy than competing models, while re-PLS additionally controls for the confounding variables (see Fig. 9 in *Supplementary Materials*). **(3)** After removing the age and gender effects, cortical thickness changes that are significantly predictive of the eight cognitive and behavioural outcomes are mainly in the temporal, frontal, and sensorimotor (see below for a discussion and Fig. 4). **(4)** The re-PLS is useful for predicting longitudinal disease progression and, particularly, seems promising to chart the disease course for subjects who change from MCI to AD over time (see Fig. 5). **(5)** The selected features and re-PLS model are generalisable and reproducible across-subjects and -cohorts (see Figs. 3 and 6).

**Fig. 2:**
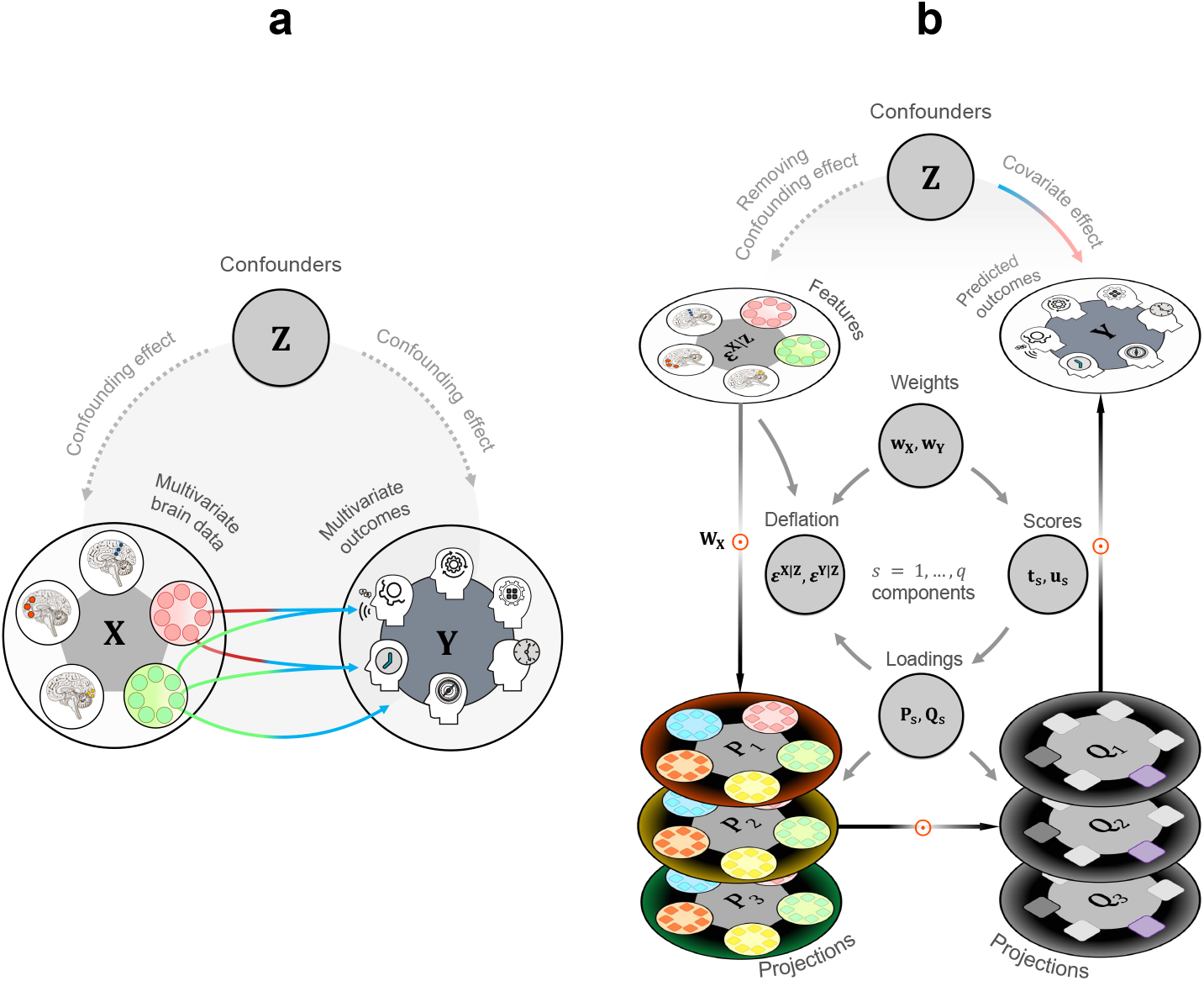
A schematic representation of Residual Partial Least Squares (re-PLS) Learning. (a) A conventional way to predict multivariate outcomes using multivariate brain features. **X** represents high-dimensional brain data, with each colored circle denoting a unique brain area and smaller circles representing cortical thickness data from that region. **Y** represents multivariate outcomes, with each icon showing a cognitive examination score (e.g., MMSE). **Z** represents confounders (e.g., age and gender) that affect both feature variables **X**, outcome variables **Y**, and the pathways between them. In classical prediction problems, one aims at looking for direct pathways between **X** and **Y** while controlling for confounding effects from **Z**. Data from the identified areas are subsequently fed via the pathways (coloured arrows) to make predictions on new subjects. (b) The Residual Partial Least Squares (re-PLS) Learning. The re-PLS begins by removing confounding effects from the confounder **Z**, isolating the residuals of both the feature matrix **X** and the outcome matrix **Y** as ***ϵ***^**X**|**Z**^ and ***ϵ***^**Y**|**Z**^, respectively. Rather than predicting directly between **X** and **Y**, re-PLS applies PLS to these residuals, iteratively learning latent components (illustrated by grey circular arrows). For each component *s*, weight vectors (**w**_**X**_, **w**_**Y**_) project data into latent scores (**t**_*s*_, **u**_*s*_) that maximize the correlation between the input and output spaces. These scores estimate loading matrices (**P** and **Q**). At each step, re-PLS performs deflation to update the residuals. The re-PLS obtains the outcome prediction through matrix multiplication (see Eq. (13)). Finally, re-PLS projects latent feature representations back to corresponding brain space to facilitate interpretation; the learned projections between the input features and the outcomes also provide clear pathways from the original features (brain-wide cortical thickness) to the multivariate disease outcomes, which are unaffected by confounders.

**Fig. 3:**
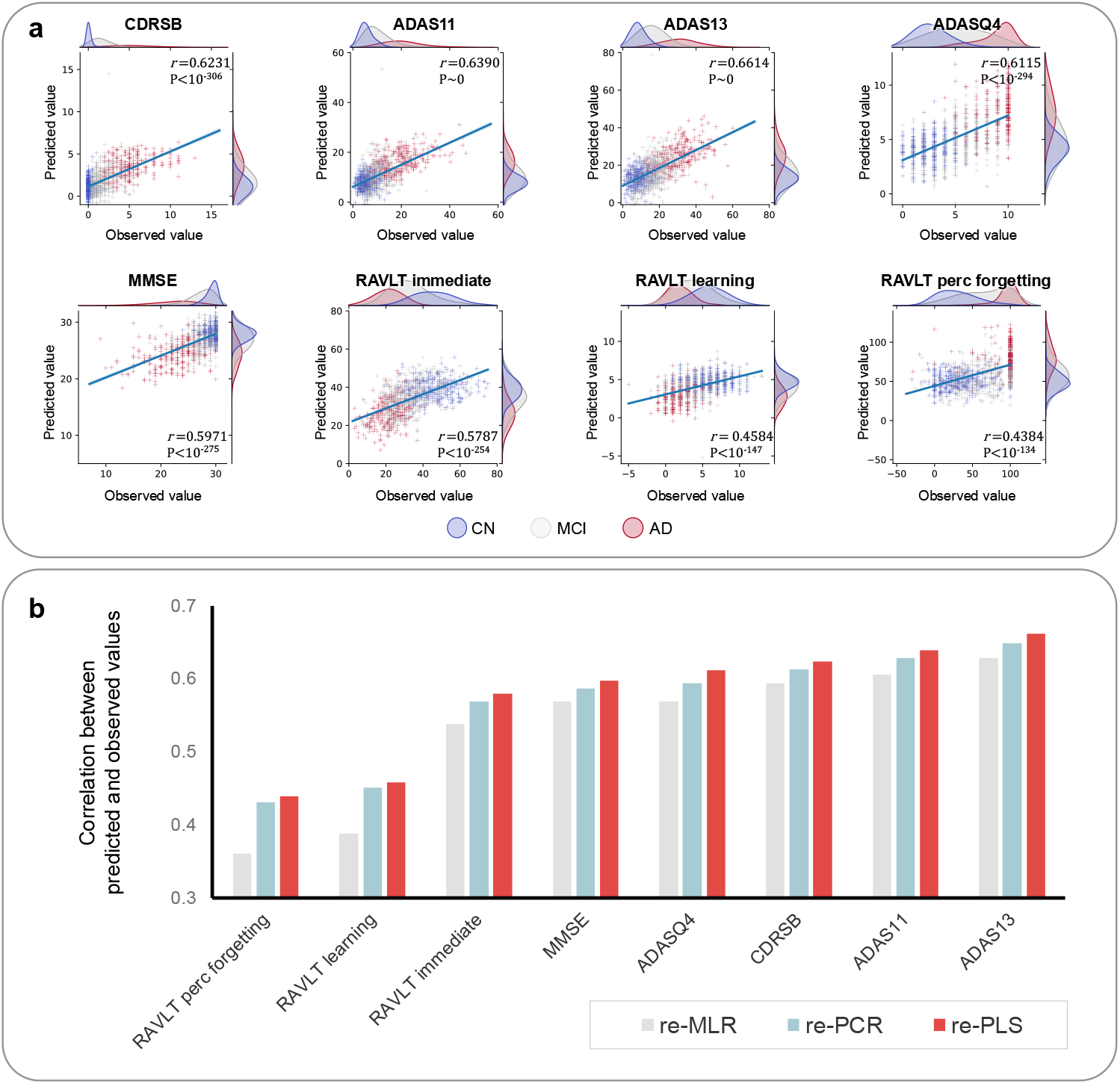
Model performance of residual Partial Least Squares (re-PLS) Learning and its performance in comparison to prominent baseline methods for predicting multivariate outcomes in previously unseen subjects. (a) Scatter plots of the predicted outcomes against the observed outcomes using re-PLS. The results are obtained by concatenating the predictions from the test set across all 10 folds of a 10-fold cross-validation (CV). CN=cognitive normal; MCI=mild cognitive impairment; AD=Alzheimer’s disease. (b) A comparison between re-PLS with two common baseline methods. Here, re-PCR refers to principal component regression with confounders controlled via residual learning, and re-MLR refers to multivariate linear regression with confounders controlled via residual learning. The plot displays the correlation coefficient between the predicted and observed outcomes, calculated from the concatenated predictions across all 10 folds. Overall, re-PLS yields the best result across eight outcomes. For both (a) and (b), only results from out-of-sample predictions were shown.

**Fig. 4:**
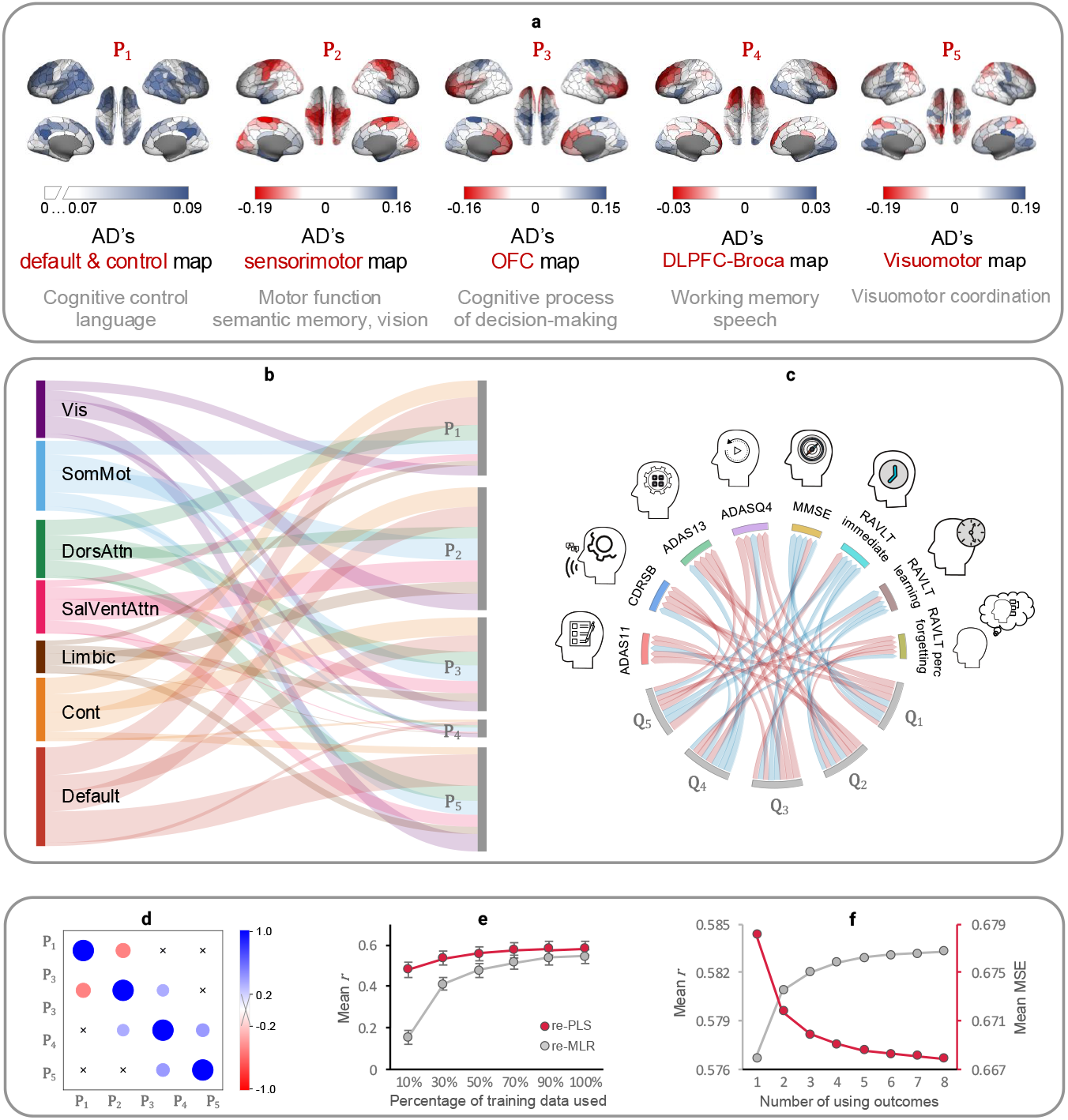
Discovering brain areas predictive of multivariate outcomes using the Residual Partial Least Squares (re-PLS) Learning. (a) The re-PLS identifies five latent brain spaces whose associated brain cortical thickness predicts multivariate outcomes in Alzheimer’s disease. The five brain spaces are discovered and verified using cross-validation. (b) The functional localisation of the five projections in terms of cortical thickness. The link between functional brain regions and the five identified projections is based on the absolute value of the weights. Each width of each projection is the sum of absolute weights; we remove non-significant lines. (c) The relationship between the outcomes’ latent spaces and the eight behavioural and cognitive outcomes through weighted connections. The width of the lines from **Q***i* (*i* = 1, …, 5) to each outcome represents the contribution of the latent representation to the outcome prediction. These weights have been normalised so that the absolute sum of each **Q** equals 1. Red lines denote negative weights, while blue lines indicate positive weights. (d) The five latent brain spaces are almost orthogonal. The size of each circle is proportional to the correlation between a pair of latent brain spaces, with “×” marking insignificant results (*P >* 0.05). (e) Sample size studies for re-PLS. During each fold, we run both re-PLS and re-MLR models using different percentages (from 10% to 100%) of the training samples and report model performance on the testing samples. (f) Prediction accuracy improves as more outcomes are included. This is an inherited property of re-PLS (through the learned projections) where each added outcome assists in the prediction of other outcomes.

**Fig. 5:**
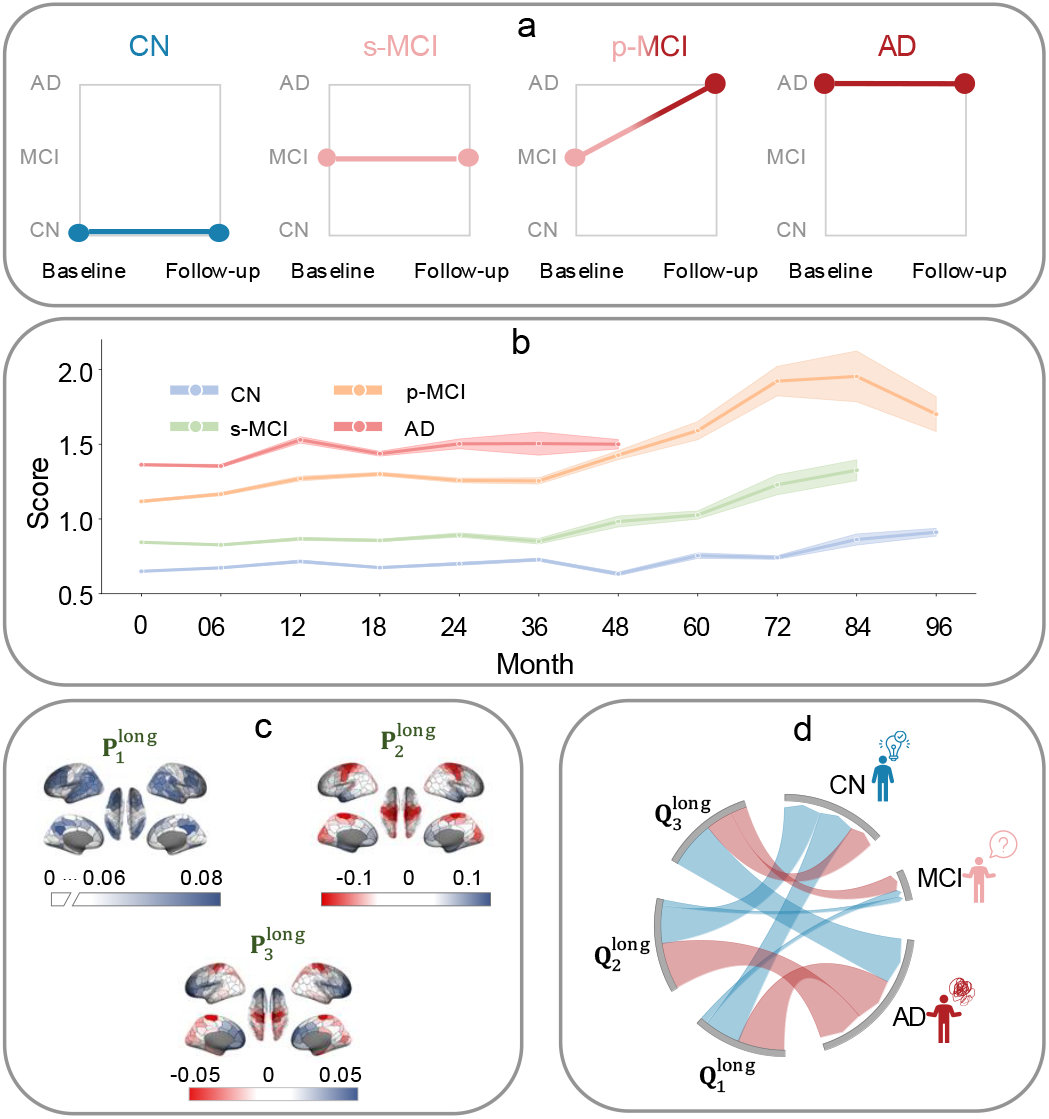
Longitudinal prediction of Alzheimer’s disease (AD). (a) Four subject groups were based on baseline and last follow-up diagnoses. CN = cognitively normal throughout, sMCI = stable MCI (MCI at baseline and follow-up), pMCI = progressive MCI (MCI at baseline, progressed to AD), and AD = diagnosed Alzheimer’s disease at both time points. Group assignment is based on the first and latest available diagnosis. (b) Longitudinal trend prediction. The longitudinal curve for each group is estimated using the predicted mean group scores for new subjects at each time point. The width of the 95% confidence bands (shaded colour) is estimated using a repeated 10-fold CV (run 100 times). In general, the predicted longitudinal severity is AD *>* pMCI *>* sMCI *>* CN. The pMCI is predicted to worsen more than other groups over time. (c) Latent brain spaces identified by re-PLS that are potentially related to longitudinal AD progression. (d) Relationship between longitudinal latent brain spaces and diagnostic outcomes. The width of the lines between three latent brain spaces and three types of diagnostic outcomes indicates the size of the association. Red lines represent negative coefficients, and blue lines represent positive coefficients. The strength of connection was estimated using the magnitude of the coefficient, quantifying the contribution each brain space makes to predicting the target outcome.

**Fig. 6:**
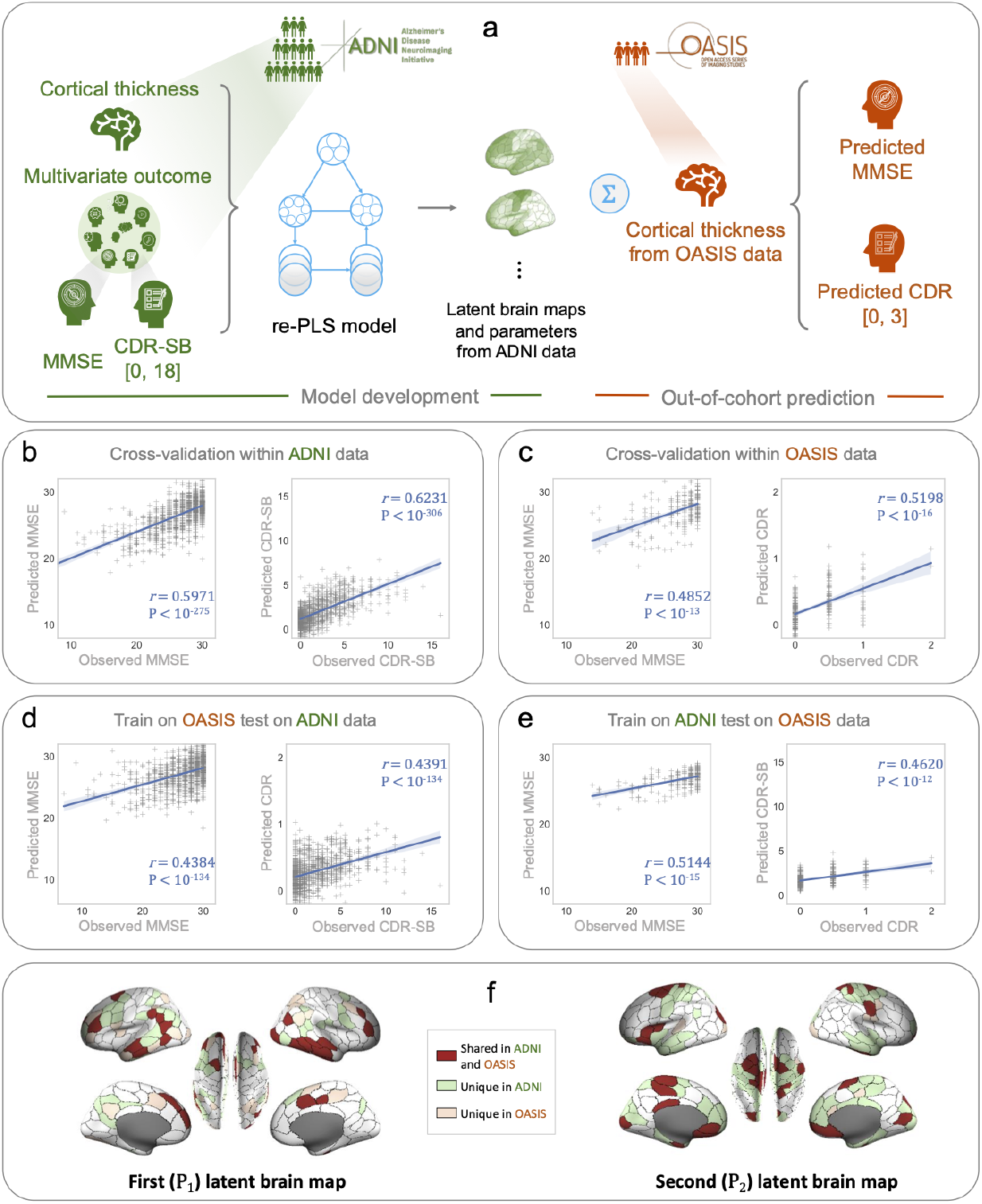
Within and cross-cohort Alzheimer’s disease (AD) prediction. (a) A schematic representation of the cross-cohort predictive model setting. From left to right, re-PLS is trained using data from the ADNI study. Extracted latent brain maps and trained parameters from ADNI data are then used, without further modelling, to predict MMSE and CDR scores in OASIS data. Note that the range of CDRSB scores is between 0 and 18, and it of CDR scores is between 0 and 3. We observe that the observed CDRSB score in ADNI is from 0 to 17, and the observed CDR score in OASIS is between 0 and 2. The setting of other within and cross-cohort analyses is similar: for within-cohort cross-validations, the training and test data are randomly selected from the same study; for ToSToL (Training on Small and Testing on Large) analysis, training data are from the smaller OASIS data, and test data are from the larger ADNI data. Out-of-sample prediction performance of: (b) Cross-subject but within-cohort analysis using ADNI data; (c) Cross-subject but within-cohort analysis using OASIS data; (d) Cross-cohort analysis (trained on OASIS data and tested on ADNI data); and (e) Cross-cohort analysis (trained on ADNI data and tested on OASIS data). (f) Visualisation of the most important and consistent brain regions for each latent map is defined by regions that are in the top 25% of absolute values and appear in 90% of 50 repetitions of a 10-fold cross-validation. Regions that are shared between ADNI and OASIS datasets are highlighted in dark red, while regions unique to the ADNI dataset are shown in light green, and those unique to the OASIS dataset are depicted in light orange.

We summarise the experimental setup in Fig. 2 and Algorithm 1. In Fig. 3, we present the model’s performance on multivariate AD outcome prediction. In Fig. 4, we identify and present the brain areas whose cortical thickness is predictive of eight cognitive-behavioural outcomes. In Fig. 5, we present the results of longitudinal AD prediction. In Fig. 6, we demonstrate the model’s ability to generalise across different cohorts (ADNI and OASIS data).

### 2.1 Cross-sectional AD assessment

We first aim to identify and separate brain regions associated with and predictive of each AD-related cognitive and behavioural outcome under a cross-sectional setting, where scans from each individual are treated as independent repeated measurements.

First, we identify anatomical regions whose cortical thickness is predictive of multivariate AD outcomes using the **P** maps (each entry of a **P** map corresponds to one brain region) (see Fig. 4). Specifically, **P**_1_ consists predominantly of areas in the prefrontal and temporal regions. **P**_2_ is largely located in the sensorimotor area with small parts in the Primary Visual Cortex (V1) and Secondary Visual Cortex (V2); it also has positive weights in parts of the temporal pole. **P**_3_ highlights the cognitive brain with a great deal of weight in the orbital prefrontal (OFC) cortex. **P**_4_ has negative weights in the dorsal lateral prefrontal cortex (DLPFC) and positive weights in parts of the Broca area. Finally, **P**_5_ contains areas with negative weights in the frontal eye fields (FEF) and BA7 (related to visuomotor coordination). We discover these brain maps via cross-subject validation using ADNI data; in Section 2.3, we further show that the patterns and the brain regions identified in **P**_1_ and **P**_2_ maps can be generalised not only across subjects but also between datasets.

Second, we investigate how the identified brain areas are distributed across functional brain regions [24]. We notice that areas in **P**_1_ are located predominantly in the default regions; areas in **P**_2_ are primarily in the sensorimotor and attention regions;areas in **P**_3_ have large representation in the default and control regions; and areas in **P**_4_ and **P**_5_ are mainly in the default and visual areas (see Fig. 4b). Furthermore, a correlation analysis between these five maps show that these projections are largely orthogonal (see Fig. 4d). This suggests that the **P** maps identify and isolate orthogonal functional brain areas that are predictive of multivariate outcomes. Based on the functional and anatomical separation (due in part to their orthogonality) of the **P** maps, we designate **P**_1_ as AD’s Default & Control map, **P**_2_ as AD’s Sensorimotor map, **P**_3_ as AD’s OFC map, **P**_4_ as AD’s DLPFC-Broca map and **P**_5_ the Visuomotor map (see Fig. 4a).

Third, the latent brain spaces (**P** maps) provide important insights about potential AD biomarkers (see Table 1). (1) Several **P** maps highlight the prominence of cortical thickness in the temporal areas (*e*.*g*., **BA20, BA21** and **BA22** in **P**_1_, and **BA38** in **P**_2_) in predicting AD-related outcomes. Previous findings suggest that the degree of atrophy in the left BA38 and BA20/21 is strongly correlated with deficits in semantic memory processing [25]. BA21, part of the middle temporal gyrus, is involved in processing language and higher-order audition processes [26]. Accumulated tau deposition in the temporal gyrus, which anatomically overlaps BA20/BA21, has been shown to be associated with clinical impairments observed in AD [27]. BA22, in the superior temporal gyrus within Wernicke’s area, is involved in the comprehension of written and spoken language. (2) Several **P** maps highlight the importance of cortical thickness in frontal regions for predicting AD-related outcomes: the prefrontal cortex (*e*.*g*., **BA9, BA10**, and **BA46**), particularly in **P**_1_ and **P**_4_; the orbitofrontal cortex (*e*.*g*., **BA10, BA11**), especially in **P**_3_, and inferior frontal gyrus (*e*.*g*., **BA44**, and **BA45**), especially in **P**_2_ and **P**_4_. BA9 (which contributes to the dorsolateral pre-frontal cortex or DLPFC) is linked with the strategic control of behaviour, including task inference, goal maintenance, inhibition, and flexible decision-making, reflecting its critical role in executive control [28, 29]. BA10 is related to working memory, episodic memory, and mentalizing [30]; it is known to support higher cognitive functions, such as task management and planning [31]. BA46 in the right hemisphere is primarily involved in spatial working memory [32, 33], whereas it in the left hemisphere is more engaged in maintaining image-based representations of objects [33]. Evidence also suggests that BA46 is significantly involved in delayed-response spatial working memory tasks [34]. BA11 is involved in decision-making, processing rewards, and encoding new information [35, 36]. BA44, which is part of Broca’s area, is involved in speech production [37]. BA45 is thought to be involved in semantic fluency [38]. (3) Parts of the parietal lobe (*e*.*g*., **BA39** and **BA40** in **P**_1_) are significantly predictive of cognitive and memory scores. This confirms previous findings that AD patients undergo cortical thickness changes in the parietal cortices [7–9]. Importantly, BA39 (angular gyrus (AG)) has been shown to correlate with longitudinal declines in verbal fluency [39]. Damage to the left BA39 may result in dyslexia or semantic aphasia [40]. BA40 (left supramarginal gyrus) is thought to be involved in reading, regarding both meaning and phonology of the words [41]. Moreover, BA39 plays a role in retrieval, particularly evident in cross-modal picture-sound pairing tasks, while BA40 plays a more limited role in sustaining retrieval [42]. (4) Our results hint at the roles sensorimotor areas (*e*.*g*., **BA4** and **BA6** in **P**_**2**_) play in predicting AD-related outcomes.

**Table 1:** Relationship between identified latent brain spaces, their anatomical correspondence, and associated functions.

| Latent brain spaces | Brodmann areas and relevant known functions |
| --- | --- |
| **P<sub>1</sub>: Default-control** | **BA9** (strategic control, action selection, and cognitive control [28, 29]), **BA10** (working memory, episodic memory and mentalizing [30]; task management and planing [31]), **BA21**, **BA22** (language and higher-order auditory processes [26]), **BA39**, **BA40** (recollective memory processes [42]), and **BA46** (spatial working memory [32]). |
| **P<sub>2</sub>: Sensorimotor** | **BA38** (semantic memory [45]), **BA17** (V1), **BA18** (V2, attentional modulation [44]), **BA4**, and **BA6**. |
| **P<sub>3</sub>: OFC** | **BA10**, **BA11** (decision making, processing rewards, and encoding new information [35, 36]), and **BA4**, **BA44**, and **BA45** (phonological and semantic processing [38]). |
| **P<sub>4</sub>: DLPFC-Broca** | **BA44**, **BA45**, **BA9**, **BA17** (V1), and **BA18** (V2). |
| **P<sub>5</sub>: Visuomotor** | **BA4**, **BA17**, **BA18**, **BA7** (visuomotor coordination [46]), and **BA8** (frontal eye fields and management of uncertainty [47]). |

Although some have argued that sensory and motor changes may precede the cognitive symptoms of AD [43], since the eight outcomes in this study measure various cognitive abilities, our findings cannot distinguish whether the changes in cortical thickness in sensory and motor areas (thus changes in sensory and motor functions) hinder movement during the examinations (thereby affecting the performance on the eight outcomes), or if they contribute, in concert with other areas, to the performance during the tests. Further research needs to independently verify this. (5) Our results suggest that cortical thickness in the visual cortex (*e*.*g*., **BA17** and **BA18** in **P**_**2**_ and **P**_**4**_) may be associated with attentional and visual memory-related word remembering. Particularly, BA18 (V2) is thought to be related to attentional modulation of visual processing [44].

Fourth, we investigated the relationship between the loadings in the latent space (**Q** loadings) and the eight behavioural and disease-related outcomes. We found that the five loadings (**Q**_1_ to **Q**_5_) exhibit overlapping associations with multiple cognitive and behavioural measures (see Fig. 4c). As each brain map **P**_*i*_ corresponds to loading **Q**_*i*_, it allows us to make interpretations both in the outcome space (through **Q**_*i*_) and regarding the brain spatial patterns (through **P**_*i*_). In particular, we observe large positive weights in **Q**_1_ and **Q**_2_ for predicting ADAS13, CDRSB, and ADAS11 scores, which measure memory, language, attention, and executive function. To understand the loadings’ neurological relevance, we enquire into their corresponding **P** maps. **Q**_1_’s corresponding **P**_1_ map (or AD’s Default & Control map) comprises lateral temporal regions, including BA20 and BA21, which is involved in semantic memory [25]. Specifically, BA20 is thought to primarily support visual association processes, while BA21 appears to be involved in audition processes and language [26]. **Q**_2_’s corresponding **P**_2_ map (or AD’s Sensorimotor map) consists of the anterior temporal lobe (BA38), which is associated with semantic memory [45]; the visual Association area (BA18/V2) exhibits enhanced effective connectivity to V5/MT during attentional modulation of motion processing [44]; and primary motor cortex (BA4) shows amyloid plaques and tau pathology in late-stage AD [48]. Next, the **Q**3 and **Q**_4_ loadings both exhibit strong predictive weights for CDRSB, ADAS13, and ADASQ4, highlighting the involvement of prefrontal and language-related regions in these cognitive outcomes. **Q**_3_’s corresponding **P**_3_ map (or AD’s OFC map) includes parts of the orbitofrontal cortex (BA10 and BA11), a region critical for decision-making, executive function, and social behaviour-domains often impaired in AD. **Q**_4_’s corresponding **P**_4_ map (or AD’s DLPFC-Broca map) comprises DLPFC and Broca’s area, essential for working memory, attention, and language production. Finally, the **Q**_5_ loading is associated with performance on ADASQ4, RAVLT immediate, and RAVLT learning, reflecting the integration of sensorimotor, attentional, and executive functions required for verbal memory and learning tasks. Relevantly, **Q**_5_’s corresponding **P**_5_ map (or AD’s Visuomotor map) includes regions involved in motor and visual processing (BA4, BA17, BA18), as well as **BA7** (visuomotor coordination [46]) and **BA8** (decision-making under uncertainty [47]).

Fifth, we investigate the impact of varying sample sizes on the performance of re-PLS compared to residual learning-aided multivariate linear regression (re-MLR). Our results suggest that, overall, re-PLS outperforms re-MLR. More specifically, it is challenging for re-MLR to perform prediction, especially when training data is small (see Fig. 4e). In comparison, re-PLS seems to deliver better overall prediction accuracy across different training data sizes and is more consistent when training data sizes vary. Additionally, re-PLS seems to require less training data to achieve optimal prediction performance. For example, to achieve comparable results using 70% of training data by re-PLS, re-MLR requires nearly 90% of the training data.

Sixth, we notice that re-PLS achieves higher prediction accuracy (both in terms of mean square error and in terms of correlation) as the number of outcomes increases (see Fig. 4f). This is possibly due to the nature of re-PLS: the hidden projections aim to maximise the associations between the inputs (cortical thickness) and outcomes (disease scores) when controlling for covariates. Thus, when making predictions, the prediction of each outcome is made by using the information of the inputs, the covariates, and the projections (which also learns information about other outcomes); although the outcomes are not all pairwise correlated, each association between two (even modestly) correlated outcomes would make one a helpful predictor of the other. Hence, the more outcomes, the better the prediction performance. Certainly, in an extreme case, when all outcomes are identical, adding additional outcomes may not improve prediction performance.

Finally, re-PLS achieves higher prediction accuracy compared to conventional linear approaches. It is likely that the lower-dimensional “nearly orthogonal” projections contain reduced noise in contrast to the original high-dimensional data. These refined (lower-dimensional) features seem to achieve more effective data representation and compression in the latent space, leading to improved prediction performance. As a result, re-PLS not only assisted neurobiological explanation via the extracted latent brain spaces but also required a smaller amount of data to achieve similar prediction performance compared to both re-PLR and re-PCR. This property may be helpful in situations with limited data but multivariate or many-to-many (high-dimensional input and multivariate outcomes) complexity.

### 2.2 Longitudinal AD assessment

As a neurodegenerative disease, AD progresses over time [3, 4]. Naturally, one would ask if it were possible to expand re-PLS to longitudinal settings to study AD progression. Longitudinal assessment is important for two reasons. First, it is beneficial to monitor and forecast disease progression to improve disease management and treatment. Second, it is useful to identify brain areas whose degenerations are related to cognitive decline over time to gain insights into how AD progresses, which brain regions contribute to disease progression over time, and, if so, to what extent.

To that end, we use re-PLS to study longitudinal AD prediction. We do so in two settings. First, we extend the use of re-PLS from cross-sectional analysis to predict AD status over time (see Fig. 5a-b). Second, we identify brain regions whose cortical thickness may be related to AD progression over time (see Fig. 5c-d). We note that we conduct longitudinal disease prediction on disease status but not on the eight outcomes. This is because conducting longitudinal multivariate disease prediction over time and across eight outcomes requires decomposing the variability into time and score space, and, therefore, requires a much larger sample size to obtain reliable results.

Throughout, we consider diagnostic outcomes made by clinicians primarily based on clinical criteria. Specifically, every subject is diagnosed with one of the statuses: CN, MCI, or AD, based on ADNI criteria. For modelling, we assign groups of 0, 1, or 2 to represent CN, MCI, and AD, respectively. In our analysis, we further group the individuals into four distinct longitudinal groups: CN, sMCI, pMCI, and AD, based on disease progression (see Fig. 5a). CN refers to individuals who were assessed as cognitively normal and maintained cognitively normal during subsequent visits. Stable Mild Cognitive Impairment (sMCI) denotes individuals who were assessed as MCIs during the first visit and continued to be diagnosed as an MCI during subsequent visits after six months. Progressive Mild Cognitive Impairment (pMCI) indicates individuals who were assessed with MCI during early visits but were diagnosed with AD during follow-up visits after six months. Lastly, AD represents individuals consistently assessed as AD throughout all visits. As some subjects have missing data at baseline, we consider their earliest scans as baseline data and arrange their later scans accordingly.

After training the longitudinal re-PLS model, we implement it to predict unseen individual subjects’ status over time. Although we grouped every subject into one of the four groups - the group information for testing subjects was not used (to avoid information leakage); rather, the (four) group information was used to colour code the testing subjects to evaluate the accuracy of the longitudinal prediction performance (see Fig. 5b). We saw that the predicted overall mean scores increased from CN, sMCI, and pMCI to AD. This agrees with the actual diagnostic outcomes. Additionally, the predicted longitudinal trend for pMCI subjects (subjects who were MCIs during early visits and later diagnosed with AD) seems to worsen noticeably more than the other groups. This is also consistent with their observed longitudinal diagnostic progression. The predicted trends for both CN and MCI groups are relatively stable,in line with their observed longitudinal diagnostic statuses, although our method predicts that both groups have a slight worsening sign after 60 months, presumptively because of a gentle cortical thinning due to ageing.

Next, we seek to unveil the brain regions whose longitudinal cortical thickness change may be potentially associated with and predictive of AD disease status over time. To that end, we extract the longitudinal latent brain spaces (**P**long maps, or the longitudinal version of the cross-sectional **P** maps), which uncover brain regions that may be associated with longitudinal disease progression over time. Specifically, we encode the disease status as a one-hot vector instead of scalars (0, 1, and 2) for the three possible outcomes (CN, MCI, and AD). Overall, we identify three longitudinal latent brain spaces. The first longitudinal map, 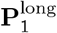, is linked to the default mode. The second longitudinal map, 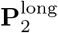, corresponds to the sensorimotor cortex, the temporal pole, and parts of the visual cortex. The third longitudinal latent brain map, 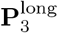, is associated with parts of the temporal, parietal, and occipital areas, and a portion of the OFC. Interestingly, all three longitudinal **P**^long^ maps (see Fig. 5c) overlap a great deal with the first three cross-sectional **P** maps (see Fig. 4a). This suggests that cortical thickness from these areas may be useful biomarkers for both cross-sectional and longitudinal AD studies.

Additionally, we investigate the **Q**^long^ loading matrix to further understand how the identified latent spaces may contribute to predicting disease status. In re-PLS framework, each 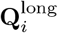 map sits along the projection of the associated 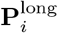 map onto the disease label space, capturing how variations in cortical thickness relate to clinical diagnosis over time. Our results show that the first two latent representations, 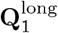 and 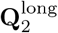, contribute significantly to the prediction of CN and AD, while 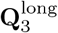 is most influential for predicting MCI. Additionally, we observe that CN status is positively associated with weights in 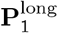 and 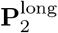 maps, but negatively associated with weights in 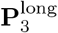 map; this pattern is reversed for AD (see Fig. 5d).

### 2.3 From cross-subject to cross-cohort AD prediction

The reproducibility of the identified latent brain maps derived from machine learning is crucial for generalising our findings, particularly for disease prediction in previously unseen subjects. In this section, we evaluate reproducibility of re-PLS and its identified neural biomarkers. We do so by expanding our cross-subject analyses using ADNI data [49] to cross-cohort studies using data from ADNI and OASIS [50]. Specifically, we consider three scenarios. First, we repeat the cross-subject analyses using ADNI and OASIS data, respectively. Second, we test the model learned and the neural biomarkers extracted from ADNI data on OASIS data. Third, we reverse the process and demonstrate the possibility of Training on Small and Testing on Large (ToSToL) data, where features extracted from smaller OASIS data are used to predict disease outcomes in much larger ADNI data.

We consider two disease outcomes, MMSE and CDR/CDRSB scores, available in both datasets. As CDRSB (used in ADNI) ranges from 0 to 18 and CDR score (used in OASIS) is between 0 and 2, to avoid data leakage and evaluate the generalisability of re-PLS and the identified neural biomarkers, we use the original scale of the CDR/CDRSB scores. Specifically, we directly trained the re-PLS using cortical thickness data and CDR-SB scoresin ADNI data, and tested the model on cortical thickness data in OASIS, and obtained predicted CDR-SB scores for OASIS samples. We then compared predicted CDR-SB scores with observed CDR scores in OASIS to assess the reproducibility of the model and neural biomarkers. We then repeat the same cross-cohort analysis, training the model on cortical thickness data and CDR scores in OASIS and evaluating it on ADNI data to obtain predicted CDR scores and compare predicted scores with observed CDR-SB scores. Additionally, to evaluate the generalizability of the latent brain maps (*e*.*g*., in Fig. 4a) during cross-cohort validation, we train the model using cortical thickness data and eight outcomes in ADNI and evaluate if the latent brain maps developed from ADNI predict MMSE and CDR in OASIS data. We note that when training the model using cortical thickness data and only MMSE and CD-SB scores in ADNI, the out-of-cohort prediction results for MMSE and CDR in OASIS improve further, and that the latent brain maps extracted using different outcomes would naturally be different. To ensure consistency, we applied the same imaging preprocessing pipeline to both datasets.

We assess the performance of the cross-cohort analyses across four scenarios (see Fig. 6). Panel (a) presents the schematic of the cross-cohort predictive modelling using re-PLS. Panels (b) and (c) portray the reproducibility of re-PLS in performing the cross-subject analysis (for ADNI and OASIS data, respectively). Panels (d)-(e) demonstrate the reproducibility of re-PLS in performing the cross-cohort analyses (trained on OASIS data and tested on ADNI data and *vice versa*). Panel (f) visualises the most important and consistent brain regions for each latent map, highlighting shared regions between the ADNI and OASIS datasets as well as those unique to each cohort. In within-cohort analyses, re-PLS generally demonstrates stronger performance compared to standard PLS. For within-OASIS analyses, the correlation between the predicted and observed MMSE scores is 0.4852 (P *<* 10^*−*13^) and the correlation between the predicted and observed CDR(SB) scores is 0.5198 (P *<* 10^*−*16^) using two latent maps learned from re-PLS. Under the same settings, PLS achieves 0.4768 (P*<* 10^*−*13^) for MMSE prediction and 0.5192 (P *<* 10^*−*16^) for CDR(SB) prediction. For within-ADNI analyses, the correlation between the predicted and observed MMSE scores is 0.5971 (P *<* 10^*−*257^), and the correlation between the predicted and observed CDR(SB) scores is 0.6231 (P *<* 10^*−*306^) using the five latent maps learned from re-PLS. See Fig. 6b and c. In comparison, PLS achieves 0.5973 (P *<* 10^*−*275^) and 0.6209 (P *<* 10^*−*304^) for MMSE and CDR(SB) predictions, respectively (see Table 1 in the *Supplementary Materials*). We highlight that, in a small number of cases here and below, where PLS achieves similar or modestly better predictive accuracy than re-PLS, the advantage of re-PLS remains in its explainability: it extracts brain regions whose pathways link features and disease outcomes that are not affected by confounders such as age and gender while maintaining predictive accuracy.

For cross-cohort analyses, our results suggest that re-PLS and the derived neural biomarkers are reproducible and generalisable between cohorts, although the prediction performance between-cohort is, as expected, slightly lower than it is within-cohort. See Fig. 6d and e. Specifically, when trained on the smaller OASIS dataset and testing on the larger ADNI dataset, re-PLS achieves 0.4384 (P *<* 10^*−*134^) and 0.4391 (P *<* 10^*−*134^) for MMSE and CDR predictions, respectively. Noticeably, when trained on the larger ADNI data and tested on the smaller OASIS dataset, re-PLS delivers good prediction performance, achieving 0.5144 (P *<* 10^*−*15^) and 0.4620 (P *<* 10^*−*12^) for MMSE and CDR predictions, respectively.

Next, we present the brain regions consistently identified across cohorts by re-PLS. In Fig. 6f, we visualise regions that are shared between ADNI and OASIS datasets, as well as cohort-specific areas. To find reproducible brain regions across cohorts, we first discover latent brain maps independently from each dataset. We then identify overlapping brain regions selected from both datasets and further evaluate their reproducibility through 50 repetitions of 10-fold cross-validation. To identify consistent regions, we threshold the areas and retain the top 25% values. We then select regions that appear consistently in 90% of the entire validation set. The first latent map (**P**_1_) highlights consistent regions in the temporal cortex, specifically BA20 and BA21, DLPFC (BA46), parts of parietal cortices (BA39), and a part of V1 on the right hemisphere, indicating a high degree of reproducibility. The second latent map (**P**_2_) also shows overlap across datasets, particularly in BA45, BA47, and parts of V2, the temporal lobe (BA20), and the sensorimotor regions (BA4/BA6). These findings suggest that the first two latent brain maps are not only predictive but also reproducible across subjects from two independent cohorts. In particular, the Default & Control (**P**_1_) map plays a dominant role in cross-cohort prediction, as evidenced by performance when trained on ADNI and tested on OASIS cohort, despite variability between cohorts. In parallel, the results also reveal dataset-specific differences. In the first latent brain map, OASIS-specific regions (in orange) are more concentrated in the occipital and parietal lobes, while ADNI-specific regions (in green) seem more concentrated in the frontal and temporal lobes. In the second latent map, OASIS-specific regions include small unique localisations in the primary visual cortex and BA43, whereas ADNI-specific regions are concentrated in the temporal, motor, and visual association areas.

Taken together, our results show that re-PLS is not only reproducible and generalisable for cross-subject (but within-cohort) analysis, but also for cross-cohort analysis: the trained model and derived neural biomarkers from one cohort predict, without further model fitting, clinically relevant outcomes in subjects from a different cohort. Additionally, our results suggest that, in addition to transferring the findings from larger datasets to smaller ones, re-PLS can be trained on smaller datasets and extrapolated to larger datasets, suggesting its potential utility in handling data size disparities for multi-centre and multi-cohort studies.

## 3 Discussion

Identifying pathways between high-dimensional multivariate brain data and multivariate, non-pairwise-correlated behavioural, cognitive, and disease outcomes is central to advancing our knowledge about how anatomical distribution and functional integration of cortical irregularities may give rise to neurodegenerative diseases. Equally, it is critical to predict disease progression that may manifest across different behavioural, cognitive, and disease domains. In this article, we develop re-PLS to (1) chart the pathways between high-dimensional multivariate brain cortical thickness data (inputs) and multivariate disease and behaviour data (outcomes); (2) simultaneously predict multiple, non-pairwise-correlated outcomes; (3) control for age and gender (confounding variables) affecting both the inputs, the outcomes, and the pathways in-between; (4) assess disease scores cross-sectionally and disease progression longitudinally; and (5) reproduce and generalise the predictive model and the selected features cross-subject and -cohort.

The re-PLS framework first obtains the residuals, unaffected by the confounders, containing information on cortical thickness and outcomes via residual learning. It then performs PLS learning between the brain data-specific and outcome-specific residuals to estimate feature weights that quantify the relationship between brain data and disease-related outcomes. The model finally uses the residuals, the confounders (now covariates), and the estimated parameters to predict multivariate outcomes in new subjects.

We first examine the method’s efficacy using data from CN subjects, individuals with MCI, and AD patients from ADNI, a multi-centre study aiming at developing biomarkers for AD [51]. Our results show that re-PLS framework is promising for identifying, separating, and estimating unique pathways between high-dimensional cortical thickness data and multivariate cognitive and behavioural scores. The identified brain regions are mainly in the temporal, frontal, and sensorimotor areas, supporting previous findings [7–9, 52–56]. Additionally, our results have provided new insights: we identify several nearly orthogonal “predictive AD biomarkers” that are jointly but differentially predictive of multivariate outcomes related to different behavioural and cognitive traits of AD. Finally, extending the model to longitudinal settings, we discover potential “longitudinal AD biomarkers” that are not only useful to explain how AD is affected spatially in the cortical areas over time but also promising to help predict longitudinal disease course and progression.

Next, to showcase the generalisability and reproducibility of re-PLS, we first perform a 10-fold CV. The model is iteratively trained on nine folds of the data and tested on the remaining fold without further model fitting (note that no subjects from the training data are in the testing set). It then iterates, training the model on nine new golds and testing it on the new remaining fold, and so on. Although our results in Fig. 4 highlight that parameters and pathways learned from the training data are helpful to predict multivariate AD outcomes in previously unseen subjects, it remains possible that the model may not capture the data variability across folds. To that end, we perform ten additional analyses with different CV settings. Specifically, we set aside *x*% (where *x* = 0, 10, 20, …, 90) of the data for an additional step of out-of-sample test and run LOOCV on (100 *− x*)% of the data; when *x* = 0, one runs LOOCV on the entire ADNI data. To avoid a (un)lucky split (*e*.*g*., the training data contains many subjects with AD and MCI, and the testing data contains many CNs), we perform stratified sampling. Taking *x* = 70 as an example, we randomly select 70% of AD subjects, 70% of the people with MCI, and 70% of CN - they form the training set, which is proportional to and representative of the entire data. The results show that the brain maps in the additional analyses, across various cross-validation settings, are generally consistent with those in Fig. 4 via 10-fold CV (see Figs. 10 to 19 in the *Supplementary Materials*). Additionally, the performance for predicting multivariate outcomes remains high and is consistent among different CV settings. Across all CV settings, the Default & Control map (**P**_1_) and Sensori-motor map (**P**_2_) are generally consistent across these 10 additional CVs. The OFC map (**P**_3_) and the DLFPF-Broca map (**P**_4_) are also consistent up to a sign (the identified key brain areas are similar with comparable weights of importance, but the signs of the weights may flip) and become increasingly stabilized as more data are used for training. The sign flip, however, does not affect interpretation and prediction. This is because (a) the method identified the same brain regions; (b) a sign flip does not affect prediction: if the **P**_*i*_ map has a sign flip, the corresponding **Q**_*i*_ also has a sign flip, thus the sign of prediction in Eq. (13) remains the same. The Visuomotor map (**P**_5_) explains the least amount of variability and is more variable across CV settings. Taken together, the additional cross-validation analyses suggest the utility of re-PLS in predicting multivariate outcomes and that the model performance and neurobiological explanation are consistent across different cross-validation mechanisms. Particularly, the consistency of the Default & Control map (**P**_1_), Sensorimotor map (**P**_2_), OFC map (**P**_3_) and DLPFC-Broca map (**P**_4_), and their convergence property as more data are used, suggest the strong plausibility of them being sensible predictive and explainable “neural biomakers” for AD. In concert, these explorations further demonstrate the generalisability and reproducibility of the method in identifying brain regions that are predictive of those non-pairwise-corrected outcomes.

Moreover, we demonstrate that the reproducibility of the neural biomarkers and re-PLS method extends to independent datasets and cross-cohort analysis using ADNI and OASIS data. These explorations show three advantages of the proposed method. First, re-PLS is not only generalisable in terms of (out-of-sample and out-of-cohort) prediction but also in terms of explanation (regarding the extracted latent biomarkers - the latent brain maps and associated parameters learned from ADNI data). Especially, the Default and Control map (**P**_1_), when directly coupled with cortical thickness data from OASIS data without further model fitting or fine-tuning, predicts MMSE and CDR scores in OASIS data. Second, the reproducibility of re-PLS in handling the ToSToL (Training on Small and Testing on Large) problem suggests that re-PLS can transfer knowledge from a smaller dataset to a larger dataset. Third, re-PLS is useful for out-of-sample or -cohort prediction, even if the scales of clinical outcomes in training and testing differ. For example, the model trained on cortical thickness data and CDR-SB scores (ranging between 0 and 18) in ADNI data predicts CDR scores (ranging between 0 and 3) in OASIS data, and *vice versa*.

There are several limitations to this study. First, the nature of the imaging and cognitive data implies that the identified pathways are associative, although our methods selected brain regions whose cortical thickness is significantly predictive of multiple cognitive outcomes (which raises association to out-of-sample and -cohort prediction). Future studies should examine whether some of the identified brain markers and pathways between the high-dimensional neural data and multiple outcomes can be raised to causal relationships. A beginning can perhaps be made by studying individuals with cortical lesions in the identified AD-related areas and examining if they exhibit AD-like behaviour and cognitive symptoms; combining re-PLS and causal inference may be helpful in this effort. Second, although re-PLS can perform longitudinal AD prediction, the algorithm was evaluated on sparse time points. This was partly due to the nature of the disease (brain structure degenerates progressively at a relatively slow pace, so it is perhaps unnecessary to have frequent assessments) and, in part, due to sparse measurements. Making a semi-continuous assessment of cognitive impairment, however, may help paint a refined, and perhaps more accurate, trajectory of the disease course, assist in monitoring symptom progression, and, for patients under treatment, evaluate the treatment efficacy more regularly and timely. Future analysis may extend re-PLS to more densely measured outcomes. Future analysis may refine patients into early and advanced AD patients and make finer forecasts. In parallel, one can apply re-PLS on MCI subjects and then follow up and apply re-PLS to data from the same subjects a few years later to study disease progression. Third, although our method unveils latent maps between brain regions and AD outcomes, the latent maps are not deep (in the sense of deep learning). One major challenge with the “deeper” models is that, while solving the many-to-many disease prediction problem, it is at present oftentimes difficult to make neurobiological sense of the identified brain areas when the weights of the (deep) hidden layers are projected on the brain space. As one of our goals here is to introduce a methodologically sound and neurobiologically meaningful method that delivers both predictive power and can identify brain areas and pathways that may shed light on neurology and neuropathology, we reserve explainable AD prediction via deep learning for future work. Fourth, the definition of AD is only based on symptoms, and the clinical diagnosis of patients only assigns them a categorical label of “AD”. Certainly, using re-PLS, we can further stratify the patients into different groups based on their continuous (non-categorical) predicted disease scores or the predicted multivariate cognitive and behaviour scores. One can even build a new, finer continuous AD total score leveraging the multivariate cognitive and behaviour scores (as different subjects have differential degeneration across those multivariate cognitive and behaviour subdomains); an example of a simple score can be a weighted sum of the predicted multivariate scores. These potentials may offer new insights about how to provide a finer prediction of the disease, but we cannot ascertain the validity using current data. Indeed, as a noticeable proportion of AD patients will end up with another diagnosis, such as FTD, LATE, PART, and vascular dementia, it is important to validate whether re-PLS can further predict AD patients into these groups. Future work can train re-PLS on subjects with FTD, LATE, PART, and vascular dementia to verify this possibility. Finally, although one of the key predictive goals in this paper is to address a many-to-many problem, re-PLS can also be used in the future to predict single outcomes (as univariate outcomes are, in essence, special cases of multivariate outcomes).

To summarise, our analyses demonstrated the possibility of identifying and isolating the many-to-many pathways between high-dimensional multivariate brain data and multiple, non-pairwise-correlated cognitive and behavioural outcomes, both cross-sectionally and longitudinally, and using the former to predict the latter in the face of confounding variables, in new subjects within the same cohort, and in subjects from a different cohort.

## 4 Methods and Materials

**Subject information**. This article uses data from the Alzheimer’s Disease Neuroimaging Initiative (ADNI) [49] and the Open Access Series of Imaging Studies (OASIS) [50].

The ADNI MRI data release includes a total of 1,196 subjects. Among them, 45 subjects are under 60 years old, 305 are in their 60s, 620 are in their 70s, and 226 are 80 or older. At baseline, 321 were CN, 28 had subjective memory complaint (SMC), 663 had mild cognitive impairment (MCI), including 234 with early mild cognitive impairment (EMCI) and 429 with late mild cognitive impairment (LMCI), and 184 were diagnosed with AD. The statuses of AD, MCI, or CN are diagnostic outcomes made by clinicians primarily based on clinical criteria (see the ADNI2 Procedures Manual at: https://adni.loni.usc.edu/wp-content/uploads/2024/02/ADNI2_Procedures_Manual_28Feb2024.pdf. During the follow-ups, 12 CNs changed to MCIs, 1 CN to AD, 2 SMCs to MCIs, and 170 MCIs to ADs. Additionally, 9 subjects with either EMCIs or LMCIs reverted to CNs, 26 subjects with SMC reverted to CNs, and 2 AD patients reverted to MCIs. For the cross-sectional study, we used data from all 1,196 subjects. For the longitudinal study, we define CN = cognitively normal, sMCI = stable MCI (a subject assessed as an MCI during the first visit and continued to be diagnosed as an MCI during subsequent visits), pMCI = progressive MCI (a subject was diagnosed as an MCI during early visits and was later diagnosed with AD), and AD = Alzheimer’s disease. We excluded 52 subjects from the longitudinal study because they were either labelled as SMC at baseline (28 subjects), converted from CN to MCI (12 subjects), or from CN to AD (1 subject), from AD to MCI (2 subjects), or from EMCI or LMCI to CN (9 subjects); they do not fall into one of the four major groups (CN, sMCI, pMCI, and AD), and their sub-sample sizes were too small to support meaningful analysis. Thus, the longitudinal study consists of 1,144 subjects, including 308 CNs, 484 sMCIs, 170 pMCIs, and 182 AD.

All participants provided written informed consent. Participants were recruited across North America and agreed to complete a variety of imaging and clinical assessments [49]. The ADNI Clinical Core manages all sites, and the Data and Publications Committee (DPC) vets all publications using ADNI data [57]. Full details regarding the initiative and the datasets are available at https://adni.loni.usc.edu/methods/documents.

This paper considers eight disease and behavioural outcomes from the Clinical Dementia Rating (CDR), the Alzheimer’s Disease Assessment Scale-Cognitive (ADAS-COG), the Mini-Mental State Examination (MMSE), and the Rey Auditory Verbal Learning Test (RAVLT). More specifically, the CDR is a score that is derived from the summation of scores from each of the six categories: Memory (M), Orientation (O), Judgment and Problem Solving (JPS), Community Affairs (CA), Home and Hobbies (HH) and Personal Care (PC). ADAS-COG assesses learning and memory, language production, comprehension, constructional praxis, ideational praxis, and orientation. It includes tasks/tests such as Word Recall, Naming, Word Recognition, Remembering Tests, Word-Finding, and Spoken Language Ability. The MMSE is a brief cognitive screening test used to assess cognitive impairment and cognitive decline. A higher score on the MMSE indicates better cognitive function, while a lower score may suggest the presence of cognitive impairment or dementia. The RAVLT assesses abilities like immediate memory, delayed recall, and recognition memory across five immediate learning trials. Further explanations regarding the scores we used in the analysis are in Table 2; full explanations of ADNI scores and procedures manual documents at https://adni.loni.usc.edu/methods/documents. We selected eight scores spanning key AD-relevant domains (memory, executive function, global cognition, and functional status) to provide comprehensive disease characterisation and demonstrate multivariate relationships across cognitive and functional dimensions.

**Table 2:**
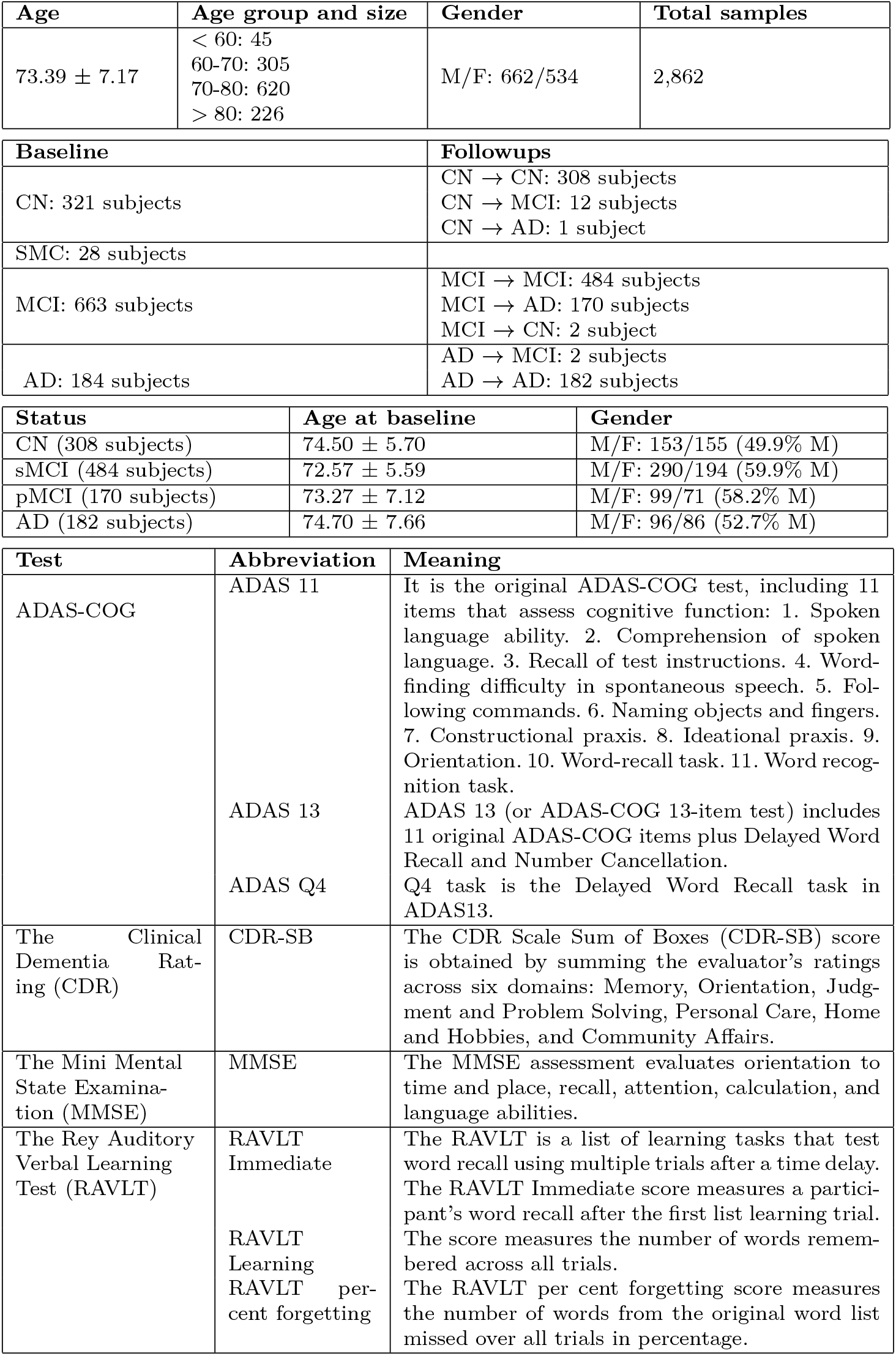
Demographic and test information for the studied sample.

| Age | Age group and size | Gender | Total samples |
| --- | --- | --- | --- |
| $73.39 \pm 7.17$ | < 60: 45<br>60-70: 305<br>70-80: 620<br>> 80: 226 | M/F: 662/534 | 2,862 |
| Baseline |  | Followups |  |
| CN: 321 subjects |  | CN → CN: 308 subjects<br>CN → MCI: 12 subjects<br>CN → AD: 1 subject |  |
| SMC: 28 subjects |  |  |  |
| MCI: 663 subjects |  | MCI → MCI: 484 subjects<br>MCI → AD: 170 subjects<br>MCI → CN: 2 subject |  |
| AD: 184 subjects |  | AD → MCI: 2 subjects<br>AD → AD: 182 subjects |  |
| Status | Age at baseline | Gender |  |
| CN (308 subjects) | $74.50 \pm 5.70$ | M/F: 153/155 (49.9% M) | |
| sMCI (484 subjects) | $72.57 \pm 5.59$ | M/F: 290/194 (59.9% M) | |
| pMCI (170 subjects) | $73.27 \pm 7.12$ | M/F: 99/71 (58.2% M) | |
| AD (182 subjects) | $74.70 \pm 7.66$ | M/F: 96/86 (52.7% M) | |
| Test | Abbreviation | Meaning |  |
| ADAS-COG | ADAS 11 | It is the original ADAS-COG test, including 11 items that assess cognitive function: 1. Spoken language ability. 2. Comprehension of spoken language. 3. Recall of test instructions. 4. Word-finding difficulty in spontaneous speech. 5. Following commands. 6. Naming objects and fingers. 7. Constructional praxis. 8. Ideational praxis. 9. Orientation. 10. Word-recall task. 11. Word recognition task. |  |
|  | ADAS 13 | ADAS 13 (or ADAS-COG 13-item test) includes 11 original ADAS-COG items plus Delayed Word Recall and Number Cancellation. |  |
|  | ADAS Q4 | Q4 task is the Delayed Word Recall task in ADAS13. |  |
| The Clinical Dementia Rating (CDR) | CDR-SB | The CDR Scale Sum of Boxes (CDR-SB) score is obtained by summing the evaluator's ratings across six domains: Memory, Orientation, Judgment and Problem Solving, Personal Care, Home and Hobbies, and Community Affairs. |  |
| The Mini Mental State Examination (MMSE) | MMSE | The MMSE assessment evaluates orientation to time and place, recall, attention, calculation, and language abilities. |  |
| The Rey Auditory Verbal Learning Test (RAVLT) | RAVLT Immediate | The RAVLT is a list of learning tasks that test word recall using multiple trials after a time delay. The RAVLT Immediate score measures a participant's word recall after the first list learning trial. |  |
|  | RAVLT Learning | The score measures the number of words remembered across all trials. |  |
|  | RAVLT per cent forgetting | The RAVLT per cent forgetting score measures the number of words from the original word list missed over all trials in percentage. |  |

In the cross-cohort setting, we use the OASIS-1 dataset [50], which includes 225 subjects with matching data to those in the ADNI: MRI images and two cognitive outcome measures: CDR (Clinical Dementia Rating) and MMSE (Mini-Mental State Examination). The MMSE scores are consistent across both datasets. The standard CDR scores range from 0 to 3, categorized as follows: 0 (no dementia, normal), 0.5 (very mild dementia, questionable), 1 (mild dementia), 2 (moderate dementia), and 3 (severe dementia); the CDR-SB (Clinical Dementia Rating Sum of Boxes) used in ADNI is more detailed, ranging from 0 (no impairment) to 18 (severe impairment).

### 4.1 Data Acquisition and Preprocessing

For the ADNI dataset, we used the preprocessed MRI images. The scans were acquired using both 1.5T and 3T with different scanner protocols in each phase (ADNI 1, ADNI 2, ADNI GO, and ADNI 3). All MRI scans in the ADNI dataset were preprocessed using the CAT12 toolbox (http://dbm.neuro.uni-jena.de/cat). We used the surface segmentation tool with default parameters to extract cortical thickness from MRI scans. For the OASIS dataset, we used the cortical thickness measurements from FreeSurfer provided by OASIS. We then used the spatial registration tool in CAT12 to map the atlas and individual brains to extract surface-based atlas maps using the Schaefer-Yeo 7 networks atlas [24] with a 200-parcel parcellation for both datasets. Secondary data analysis, including re-PLS, was conducted using a customised Python package available at https://github.com/thanhvd18/rePLS.

#### Algorithm 1

The Residual Partial Least Squares Learning

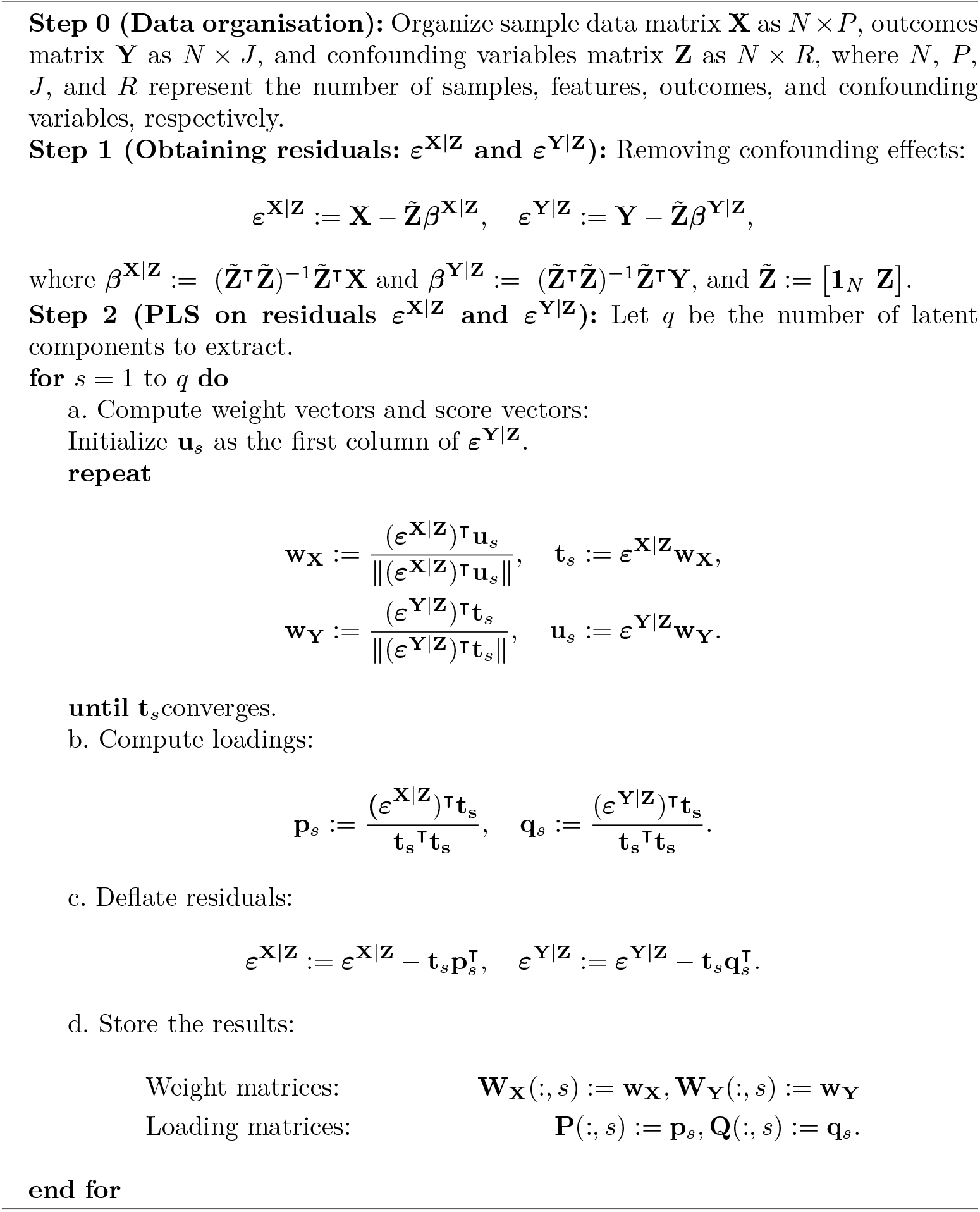

#### Notations and data organizations

We begin by defining the notations used throughout this article. Let **X** ∈ ℝ^*N ×P*^represents the input data with *N* subjects and *P* features. Let **Y** ∈ ℝ^*N×J*^ and **Z** ∈ ℝ^*N ×R*^ be the outcome and confounder matrices, respectively. Each subject *i* (1 ≤ *i ≤ N*), we define *y*_*ij*_ and *z*_*ir*_ as the *j*^*th*^ outcome and the *r*^*th*^ confounding variable, respectively, for 1 *≤ j ≤ J*, and, 1 *≤ r* ≤ *R*. The dataset is partitioned into training and test sets of sizes *N*_train_ and *N*_test_ respectively (*N* = *N*_train_ + *N*_test_). The corresponding subsets of data are written as 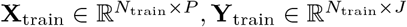, and 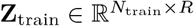, with similar definitions for the test set.

#### Residual Partial Least Squares (re-PLS) Learning

We outline re-PLS framework, which adjusts for confounding variables before performing partial least squares regression. The process begins by computing residuals of the predictor and outcome matrices with respect to the confounders **Z**. Specifically, during training, we define:

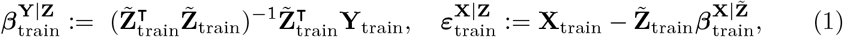

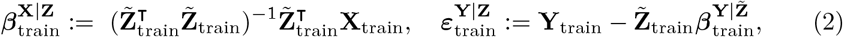

where 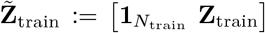 and **1***N*_train_ used to add an intercept term in linear regression. This step makes sure 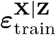 and 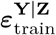are both zero-centered. For a test subject, we compute:

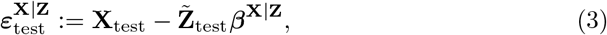

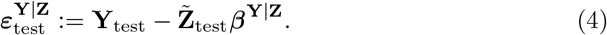

Note that the confounding structure **Z** is effectively removed from both **X** and **Y** through linear regression, so the confounders no longer influence the residuals. To make predictions on unseen data, we use:

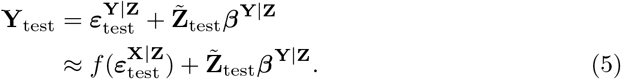

Here, the function *f* (*·*) represents the mapping from residualised covariates to resid-ualised outcomes. To learn this relationship, we perform PLS on residuals 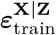 and 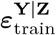, a process which we term *Residual PLS Learning* (re-PLS). The two key points of performing re-PLS are:

1. After removing the confounding effect, the residuals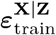 and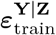 to provide better insights about the potential relationship (see Section 1 of the *Supplementary Materials*) between the multivariate features **X**_train_ and outcomes **Y**_train_ (compared to the case when confounder effect exists), as the residuals still contain information about **X**_train_ and **Y**_train_ but are independent of **Z**_train_.
2. After removing the effect of **Z**_train_ on **X**_train_, we consider 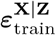as the new, trans-formed input variable (or transformed features), and the initial confounding effect of **Z**_train_ on **Y**_train_ now becomes a covariate effect (note that 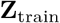 affects **Y**_train_, 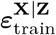 affects **Y**_train_, but **Z**_train_ does not have any effect on 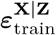). This observation is valuable for performing out-of-sample prediction.

In the following, we outline the second key point of rePLS learning. We project data into latent space instead of directly regressing the outcome on the input. We then learn regression coefficients in this latent space and transform the predictions back to the original variable space.

Each component (denoted as *s*) is learned at one time. The first step is to learn weight vectors (**w**_**X**_ and **w**_**Y**_) that maximize the covariance between score vectors:

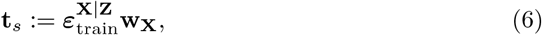

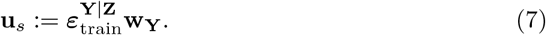

For simplicity, in the remaining part of the paper, we use notation without “train” to denote data used during training or parameters estimated from training data (for example, 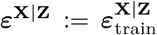and 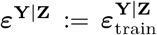). The score vectors are updated itera-tively by alternating updates until convergence. Then the score vector **t**_*s*_ captures the direction in ***ε***^**X**|**Z**^ that has the highest covariance with ***ε***^**Y**|**Z**^. The loading matrices are obtained by regressing ***ε***^**X**|**Z**^ and ***ε***^**Y**|**Z**^, respectively, on the score vector **t**_*s*_.

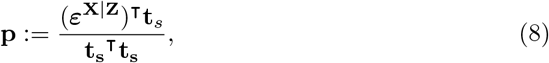

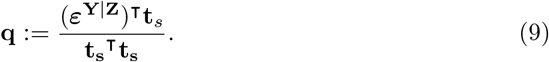

The data matrices ***ε***^**X**|**Z**^ and ***ε***^**Y**|**Z**^ are deflated and then utilised to learn the next component.

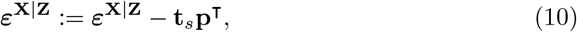

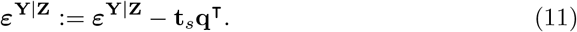

After computing all components, we store the weight and loading vectors in the matrices **w**_**X**_ ∈ ℝ^*P ×I*^, **w**_**Y**_ ∈ ℝ^*P ×J*^, **P** ∈ ^*P ×I*^, and **Q** ∈ ℝ^*P ×J*^ to make the final prediction. Since the columns of **P** are generally not orthogonal, the latent score matrix **T** cannot be directly recovered from ***ε***^**X**|**Z**^ using **P**. Instead, we use the following relation:

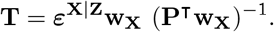

This gives us the following form for predicting residualised outcomes from residualised covariates:

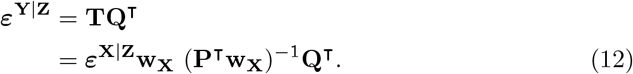

We outline the entire procedure for computing these components in Algorithm 1 and provide simulation studies and model comparisons with other methods inSection 3 of the *Supplementary Materials*.

#### Predict multivariate outcomes in new subjects

Consider new subjects with feature data **X**_test_ and confounders **Z**_test_. The predicted outcome **Y**_test_ for these new subjects without additional model fitting is given by:

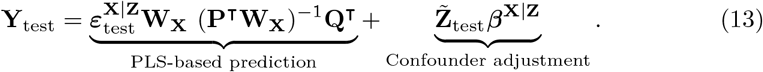

Note that after removing the confounding effect of **Z**_test_ on **X**_test_, the residuals 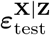 are no longer affected by **Z**_test_. Therefore, the effect on the outcomes is now a covariate effect, namely the second part in Eq. (13).

## Code availability

The re-PLS Python source code is available at https://github.com/thanhvd18/rePLS.

## Data availability

The ADNI data used in the preparation of this article were obtained from the Alzheimer’s Disease Neuroimaging Initiative database and are available through an application at https://adni.loni.usc.edu/data-samples/adni-data/#AccessData. The OASIS data used in this article were provided by OASIS-1 and are available through an application at https://sites.wustl.edu/oasisbrains/home/access.

## Author contributions

D.T.V. and O.Y.C. conceptualised the study. D.T.V. and O.Y.C. developed the methods. D.T.V. wrote the code, performed data processing and analysis, and wrote and maintained the re-PLS package. D.T.V. obtained all the results and figures in the manuscript. N.L.T. and O.Y.C. provided support and guidance for D.T.V. H.C., G.A., P.R., and G.N. provided neurobiological support. C.S.D., J.S.B., G.E.B., H.P., V.D.N., H.S., and H.Z. provided statistical and machine learning support. X.H., B.Z., W.H., X.W., and M.M. provided insights into disease prediction. Y.M. and G.P. provided clinical and medical insights. O.Y.C., D.T.V., and D.C.C. wrote the manuscript, with comments from all other authors.

## Acknowledgements

The ADNI data for this project were funded by the Alzheimer’s Disease Neuroimaging Initiative (ADNI) (National Institutes of Health Grant U01 AG024904) and the Department of Defence ADNI (Department of Defence award number W81XWH-12-2-0012). The investigators within the ADNI contributed to the design and implementation of ADNI and/or provided data, but they did not take part in the analysis or preparation of this report. A full listing of ADNI investigators is available at: http://adni.loni.usc.edu/wp-content/uploads/how_to_apply/ADNI_Acknowledgement_List_pdf. ADNI is funded by the National Institute on Aging, the National Institute of Biomedical Imaging and Bioengineering, and through generous contributions from the following: AbbVie, Alzheimer’s Association; Alzheimer’s Drug Discovery Foundation; Araclon Biotech; BioClinica, Inc.; Biogen; Bristol-Myers Squibb Company; CereSpir, Inc.; Cogstate; Eisai Inc.; Elan Pharmaceuticals, Inc.; Eli Lilly and Company; EuroImmun; F. Hoffmann-La Roche Ltd and its affiliated company Genentech, Inc.; Fujirebio; GE Healthcare; IXICO Ltd.; Janssen Alzheimer Immunotherapy Research & Development, LLC.; Johnson & Johnson Pharmaceutical Research & Development LLC.; Lumosity; Lundbeck; Merck & Co., Inc.; Meso Scale Diagnostics, LLC.; NeuroRx Research; Neurotrack Technologies; Novartis Pharmaceuticals Corporation; Pfizer Inc.; Piramal Imaging; Servier; Takeda Pharmaceutical Company; and Transition Therapeutics. The Canadian Institutes of Health Research provides funds to support ADNI clinical sites in Canada. Private sector contributions are facilitated by the Foundation for the National Institutes of Health (www.fnih.org). The grantee organisation is the Northern California Institute for Research and Education, and the study is coordinated by the Alzheimer’s Therapeutic Research Institute at the University of Southern California. The Laboratory disseminates ADNI data for Neuro Imaging at the University of Southern California.

The OASIS data used in this article were provided by OASIS-1: Cross-Sectional: Principal Investigators: D. Marcus, R, Buckner, J, Csernansky, and J. Morris; P50 AG05681, P01 AG03991, P01 AG026276, R01 AG021910, P20 MH071616, U24 RR021382.

